# Positive relation between arcuate fasciculus white matter fiber structure and severity of auditory hallucinations: A DTI tractography study

**DOI:** 10.1101/784942

**Authors:** Liv E. Falkenberg, René Westerhausen, Erik Johnsen, Rune Kroken, Else-Marie Løberg, Justyna Beresniewicz, Katarzyna Kazimierczak, Kristiina Kompus, Lars Ersland, Kenneth Hugdahl

## Abstract

The arcuate fasciculus (AF) has been implicated in the pathology behind schizophrenia and auditory verbal hallucinations (AVHs). White matter tracts forming the arcuate fasciculus can be quantified and visualized using diffusion tensor imaging (DTI) tractography. Although there have been a number of studies on this topic, the results have been conflicting. Studying the underlying white matter structure of the AF could shed light on functional connectivity between temporal and frontal language areas in AVHs. The participants were 66 patients with a schizophrenia diagnosis, where AVHs were defined from the Positive and Negative Syndrome Scale (PANSS), and compared with a healthy control group. DTI was performed on a 3T MR scanner, and tensor estimation was done using deterministic streamline tractography. Statistical analysis of the data showed significantly longer tracts along the AF in patients with severe and frequent AVHs, as well as an overall significant asymmetry with longer fibers on the left side. In addition, there were significant positive correlations between PANSS scores and tract length, tract volume, and number of streamlines for the posterior AF segment on the left side. It is concluded that the present structural results complement previous functional findings of fronto-temporal connectivity in AVH patients.

## Introduction

One of the most salient symptom in schizophrenia and psychosis spectrum disorders is auditory verbal hallucinations (AVHs) (Sartorius et al., 1986; Parnas, 2013; Ford et al, 2014). AVHs are typically defined as the experience of hearing a “voice” in the absence of a corresponding external auditory source to explain the experience (Waters et al., 2006; Aleman & Larøi, 2008; Hugdahl, 2015; Pienkos et al., 2019 for a conceptual review). Looking for neuronal correlates of AVHs, several meta-analyses have implicated the language regions in the temporal and frontal lobes, particularly in the left hemisphere (Curcic-Blake et al., 2017; Jardri et al., 2011; Kompus et al., 2011, Kühn & Gallinat, 2010; Sommer et al., 2012). Functional connectivity studies, using various statistical approaches such as seed-voxel correlations and independent component analysis, have added to the earlier imaging studies by showing altered connectivity between auditory and language regions (see Alderson-Day et al., 2015 for a review). The existing literature is however inconclusive when it comes to the direction of the alterations in AVH patients, with some studies showing an increase in specific and global connectivity (e.g. Diederen et al., 2013; van Lutterwald et al., 2014; Chang et al., 2017; Zhou et al., 2020), while other studies have shown a decrease in connectivity (Sommer et al., 2012; Clos et al., 2014). Whether AVH causes increased or decreased functional grey matter connectivity is therefore an unresolved issue (see Scheinost et al., 2019, and Alderson-Day et al., 2014).

Functional fMRI connectivity is based on the statistical covariation of the BOLD signal between nodes or areas (disregarding the special case of effective connectivity) without any reference to the underlying neuroanatomy, or whether an increase or decrease in functional connectivity has a correspondence in structural, white matter, connectivity. Hugdahl (2017) labelled this lack of “convergence of evidence”, by which was meant that for a finding at one level of explanation to be valid, there should be converging evidence obtained from a lower, or different, level of explanation (see also Hugdahl & Sommer, 2018). A converging approach would therefore be to look for white matter connectivity between brain anatomical regions implicated in functional connectivity studies. Two brain regions that are often implicated in functional neuroimaging studies of AVHs are the superior temporal gyrus, including the transverse, Heschl’s, gyrus and the planum temporale on the one hand, and the inferior frontal, triangular, gyrus, on the other hand (Sommer et al., 2010; Kompus et al., 2011, Jardri et al., 2011). Functionally, these regions constitute the classic Wernicke and Broca regions for language perception and production, respectively, and these regions have over the years been associated with AVHs (Woodruff, 1997; Hubl, 2008; Loijestin et al., 2013; Curcic-Blake et al., 2017; Alderson-Day et al., 2015 for reviews). The arcuate fasciculus (AF) (see Figure 1) is a massive fiber bundle which runs longitudinally along the lateral anterior-posterior axis, connecting the Wernicke, speech perception area in the superior temporal gyrus with the Broca speech production area in the inferior frontal gyrus on the left side, with a corresponding extension on the lateral side of the right side. The AF consists of both short and long fibers, the short fibers connecting areas within a region, while the long fibers connect between regions (Catani et al., 2011; Catani & Thiebaut de Schotten, 2008).

**Figure 1:**
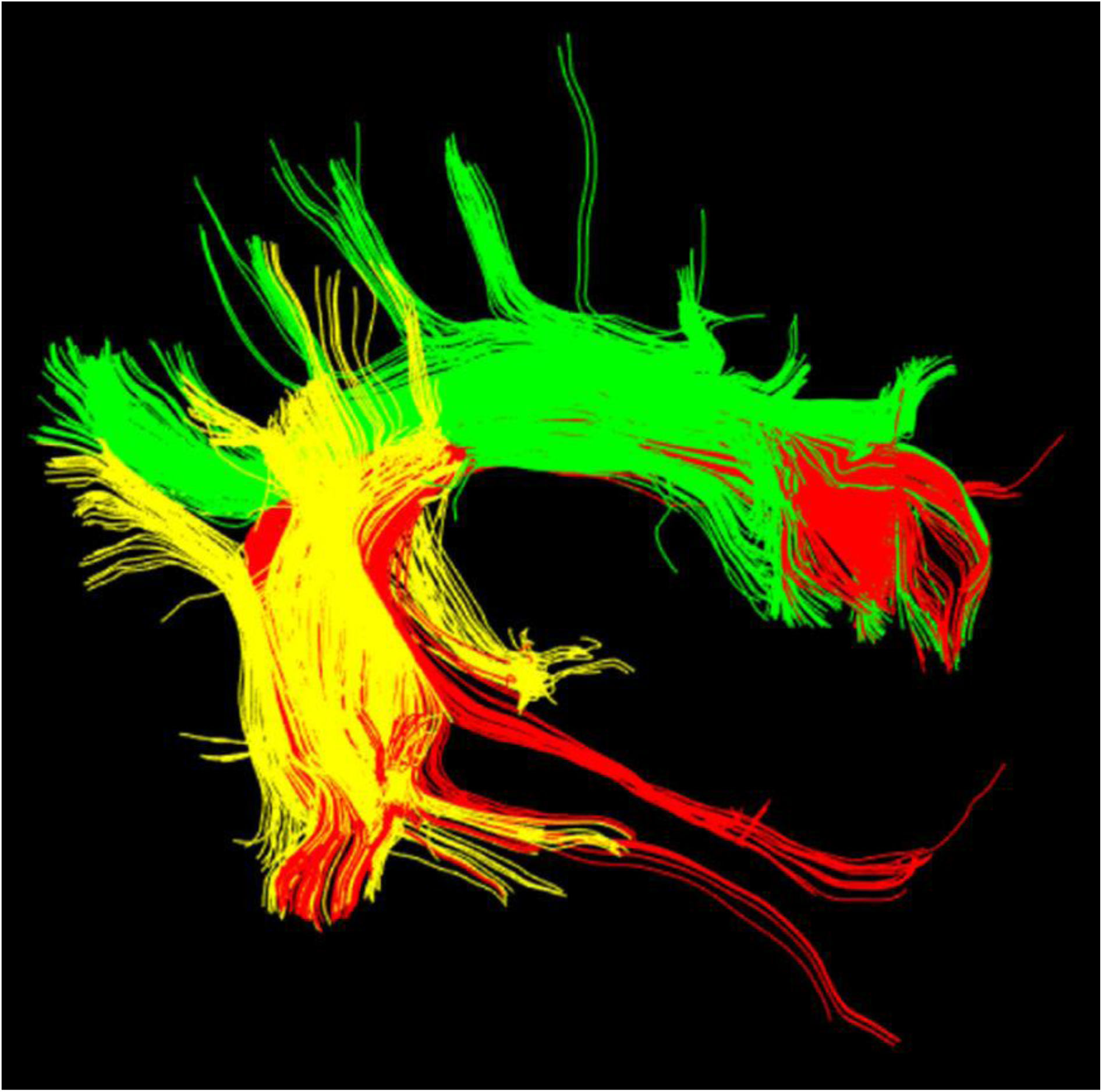
**T**he three segments of the arcuate fasciculus (AF): Anterior (green), long (red), and posterior (yellow). Adapted after Catani et al. (2011).

The AF is suggested to be important for language processing, connecting temporal auditory areas with inferior frontal/precentral and inferior parietal regions. Using diffusion tensor imaging (DTI), Fernandez-Miranda et al. (2015) found a clear asymmetry across the hemispheres for the AF with the left AF fiber volume significantly larger than the homologous right volume (see also Ocklenburg et al, 2013 for similar findings).

The AF has also been implicated in the pathology behind schizophrenia and AVHs, most commonly by findings of disruptions of white matter integrity. In a meta-analysis, Geoffroy et al. (2014) found reduced fractional anisotropy (FA) values, quantified from diffusion tensor imaging (DTI) data, in schizophrenia patients with frequent AVHs, reflecting abnormality of white matter organization along the AF extension. Although only five studies were included in the meta-analysis, the results nevertheless pointed to the importance of relating findings of functional connectivity between temporal and frontal language areas to the underlying white matter structure. Previous studies have also found an increase in white matter volume in the regions connected by the AF (Mitelman et al., 2007), and that greater overall white matter volume predict higher positive symptom scores, including hallucination scores (Cahn et al., 2006).

In the present study we sought to expand previous findings of altered connectivity in patients with severe and frequent AVHs (Scheinost et al., 2019, and Alderson-Day et al., 2014 for overviews), by acquiring data from diffusion tractography to converge evidence from functional to structural levels of explanation (cf. Hugdahl & Sommer, 2018). We used the sub-division of AF fibers and the nomenclature from Catani et al. (2011), divided the AF into its anterior, long and posterior fiber segments (see Figure 1 for illustration), and analyzed each segment for number of tracts (Tract#), tract length (TractL), and tract volume (TractV), comparing the fibers in the right and left hemisphere, for schizophrenia patients with and without auditory hallucinations (as defined operationally from the Positive and Negative Syndrome Scale, PANSS, interview questionnaire). We hypothesized that if an increase in AF fiber connectivity, either increasing fiber length, volume or number of fibers, this would strengthen previous functional connectivity studies of increase in functional connectivity between temporal and fronto-parietal areas, while the opposite would be expected if a decrease in AF fiber connectivity was found. Second, we hypothesized that any effects would be more pronounced on the left side, considering previous studies of language asymmetry being constrained to the left hemisphere.

## Methods

### Subjects

The subjects were 66 patients with a diagnosis of schizophrenia (SZ), according to the ICD-10 or DSM-IV (American Psychiatric Association, 2008) diagnostic manual systems, 40 males and 26 females, with average age for the whole group 27.02 years (SD 9.18), and mean duration of illness of 4.32 (SD 5.01) years. All the patients used either first- or second-generation antipsychotics. The patients were compared with a healthy control (HC) group consisting of 76 subjects, 47 males and 29 females, with an average age of 28.69 (SD 7.44) years (for some of the analyses the control group was reduced to 68 and the patient group to 63 subjects due to missing data). There were 54 right-handers in the SZ group, and 67 right-handers in the HC group. The SZ group was further divided into sub-groups depending on severity and frequency of auditory hallucinations as defined from the “hallucinatory behavior” item (P3) from the PANSS (Kay et al., 1987). Using the criterion of a score equal to or larger than “4” on the P3 item split the SZ group into a AVH+ sub-group (n = 30, 15 males, 15 females), and AVH- (n = 36, 25 males, 11 females). The patient data were pooled from two studies, each approved by the Regional Committee for Medical Research Ethics in Western Norway (REK Vest), #2010/334 and #2016/800, respectively, and by the Norwegian Social Science Data Service (NSD) #15733/PB.

### Diffusion weighted MR imaging

Diffusion weighted imaging was performed on a 3T GE Signa HDx MR system with the following scanning parameters; TE = 89 ms, flip angle = 90°, TR = 14000 ms, FoV = 220 mm, matrix = 128 x 128, Slice thickness, 2.4 mm, with total scan time 8.38 min. Diffusion sensitizing gradients were applied in 30 directions, with a weighing factor of b = 1000 s/mm^2^, with 6 b0 images as reference, and a reconstructed voxel size of 1.72 x 1.72 x 2.24 mm. There was a scanner upgrade in about the middle of the project period, upgrading from the GE Signa HDx to the GE Discovery 750 system.

### Tensor estimation and tractography

Data quality assessment, preprocessing, tensor estimation and tractography were done using the ExploreDTI v. 4.8.4 software (Leemans et al. 2009). First, diffusion weighted images were corrected for movement (including b-matrix rotation) and visually inspected for artefacts. Following voxel-wise tensor estimation, whole brain deterministic streamline tractography (tracking threshold FA = 0.2, angle threshold = 30°) was conducted. Then, for each individual and hemisphere, the anterior, long, and posterior segments of the AF were segmented according to landmarks described by Catani et al. (2011). Manually placed “way masks” were drawn in the coronal and axial plane, with ‘exclusion-masks’ eliminating fibres from the long segment in the anterior and posterior segment. Tractography was done in native space. Mean tract volume (TractV), tract number (Tract#), and tract length (TractL) were extracted as macrostructural measures.

### Statistical analyses

A first analysis was set up as a multifactorial general linear mixed model (GLM) analysis of variance (ANOVA), separately for the TractL, Tract#, and TtractV measures, with Segments (anterior, long, and posterior) and Hemisphere (left/right) as within-subject factors, and Group (AVH+, AVH-, HC) as between-subjects factor, and with Age and Sex as covariates. Significant main and interaction F-values were Geisser-Greenhouse epsilon-corrected when appropriate for inflated degrees of freedom. Fisher’s LSD test was used for follow-up tests of significant main- and interaction-effects, with an alpha level set to p <.05. A second analysis was to correlate values for Tract#, TractL, and TractV with scores for the hallucinations item (P3), which would then provide a quantification of the association between AF white matter structure integrity and severity and frequency of AVH. Spearman’s rank correlations were applied because of the restricted range of the P3 values from 1 to 7, in practice from 1-6 since patients are seldom scored by clinicians as extreme hallucinators as a score of “7” would require.

## Results

### Tract length (TractL)

Figure 2 illustrates the long fibers in three representative subjects, one from each group of AVH-, AVH+ and HC.

**Figure 2:**
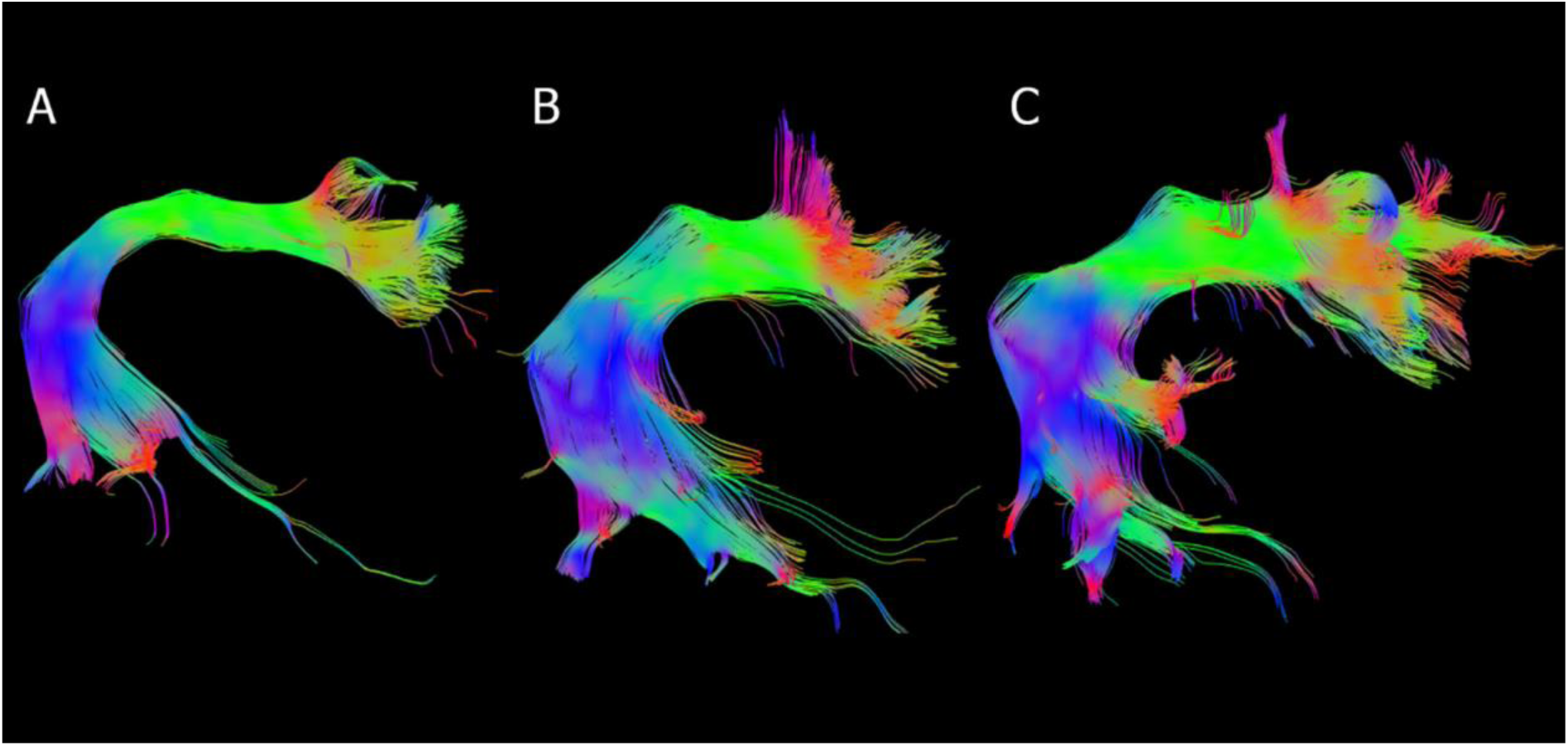
Representative long, direct segments of the left arcuate fasciculus from A) a healthy control, B) a schizophrenia patient without AVHs (PANSS P3=1), and C) a schizophrenia patient with frequent AVHs (PANSS P3=5). Note: Radiological convention applied. AVHs, auditory verbal hallucinations; PANSS, positive and negative syndrome scale; P3, positive item number 3 – hallucinatory behavior.

The ANOVA showed a significant main-effect of Segment, F(2,250, G-G corrected df: 1.494, 186.80) = 145.029, p < .0001, partial eta^2^ = .537, and of the interaction of Segment x Group, F(4, 250, G-G corrected df: 2.989, 186.83) = 3.020, p = .018, partial eta^2^ = .046. In addition, the Group effect was marginally significant, F (2,125) = 2.999, p = .053, partial eta^2^ = .045. The interaction-effect was followed-up with the LSD-test, comparing the three groups for each segment separately. For left hemisphere comparisons, this showed significant differences between the HC group and the AVH+ group (p = .058) for the long segment, and a corresponding trend for the difference between the AVH+ and AVH-groups (p = .085). For the right hemisphere, the corresponding comparisons for the right hemisphere were p = .005 for the comparison HC versus AVH-group, and p = .0003 for the comparison HC versus AVH+, while the comparison of AVH-versus AVH+ was not significant, p = .362 (see Figure 3). There was in addition a significant tract length asymmetry for the HC group with longer tracts on the left side, p = .0001, and a trend for a similar asymmetry for the AVH+ group (p = .074 (see Figure 3).

**Figure 3:**
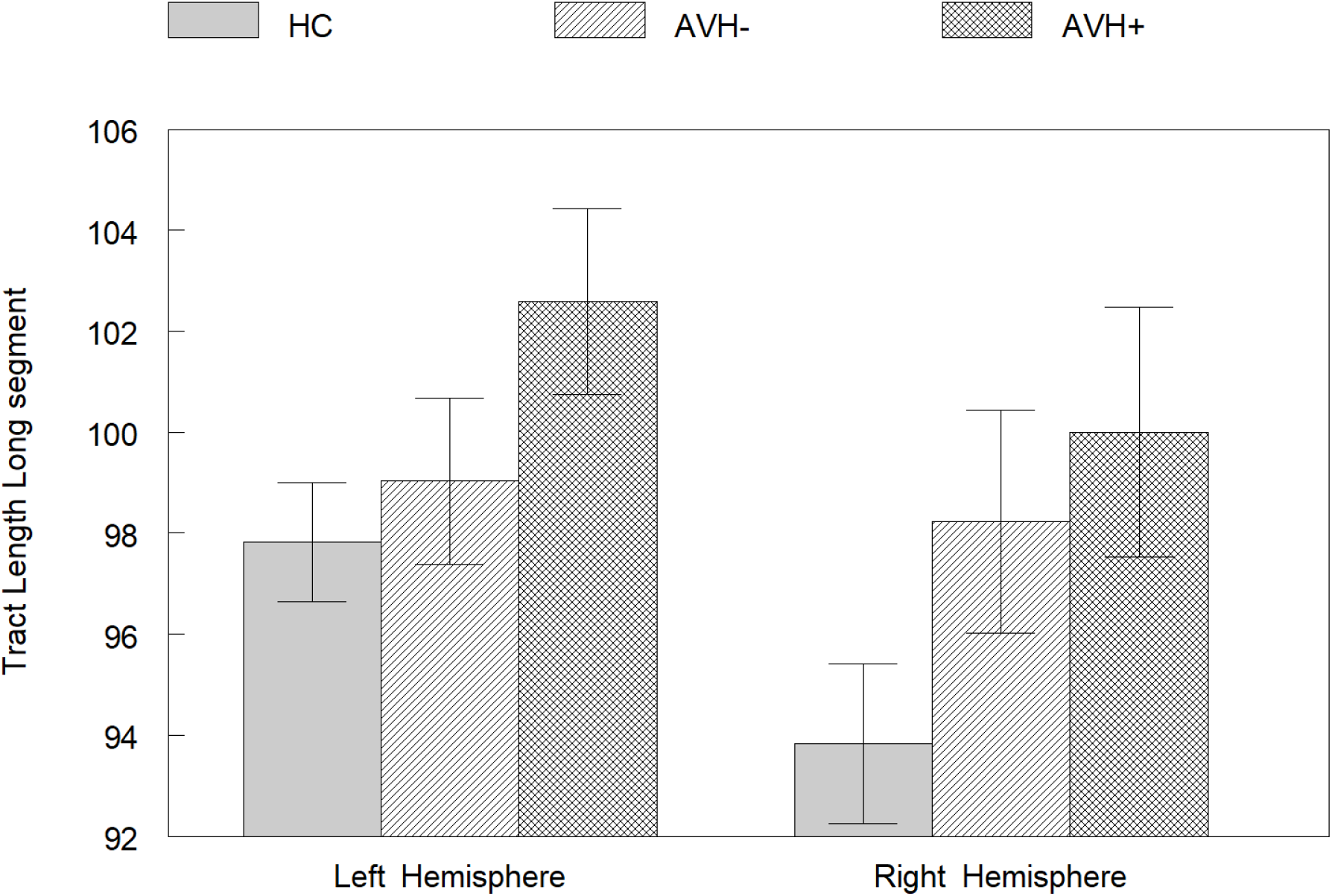
Means and SE for the three groups HC (healthy controls), AVH-(no-hallucinations), AVH+ (hallucinations) group for the Tract Length long tract segment data.

### Number of tracts (Tract#)

There were no significant main- or interaction-effects involving Groups for number of tracts.

### Tract volume (TractV)

There were no significant main- or interaction-effects involving Groups for tract volume.

### Correlations

Spearman correlation coefficients for the correlations between the various tracts and segments on the one hand, and PANSS P3 score on the other hand, are seen in Table 1.

**Table 1:** Spearman correlation coefficients for correlations between PANSS P3 scores and number of tracts (Tract#), tract length (TractL), and tract volume (TractV), separately for the left and right hemisphere

|  | PANSS P3 | PANSS P3 |
| --- | --- | --- |
|  | Left Hemisphere | Right hemisphere |
| Tract#_Ant | -0,0902 | -0,103572 |
| TractL_Ant | -0,0616 | 0,040424 |
| TractV_Ant | -0,1339 | -0,104622 |
| Tract#_Long | 0,0637 | 0,176181 |
| TractL_Long | 0,1618 | 0,051332 |
| TractV_Long | -0,0054 | 0,203135 |
| Tract#_Post | 0,3300 | 0,176181 |
| TractL_Post | 0,2720 | 0,051332 |
| TractV_Post | 0,3470 | 0,203135 |

The correlations were focused on significant relationships between severity and frequency of AVH (as measured from the PANSS P3 score) and tract segments (as measured from the procedure described in the Methods section). As seen in Table 1, there were significant positive correlations (p < .05) for the posterior segment of the tract length in the left hemisphere. Figure 4 shows the corresponding scatter-plots, separately for the left and right hemisphere.

**Figure 4:**
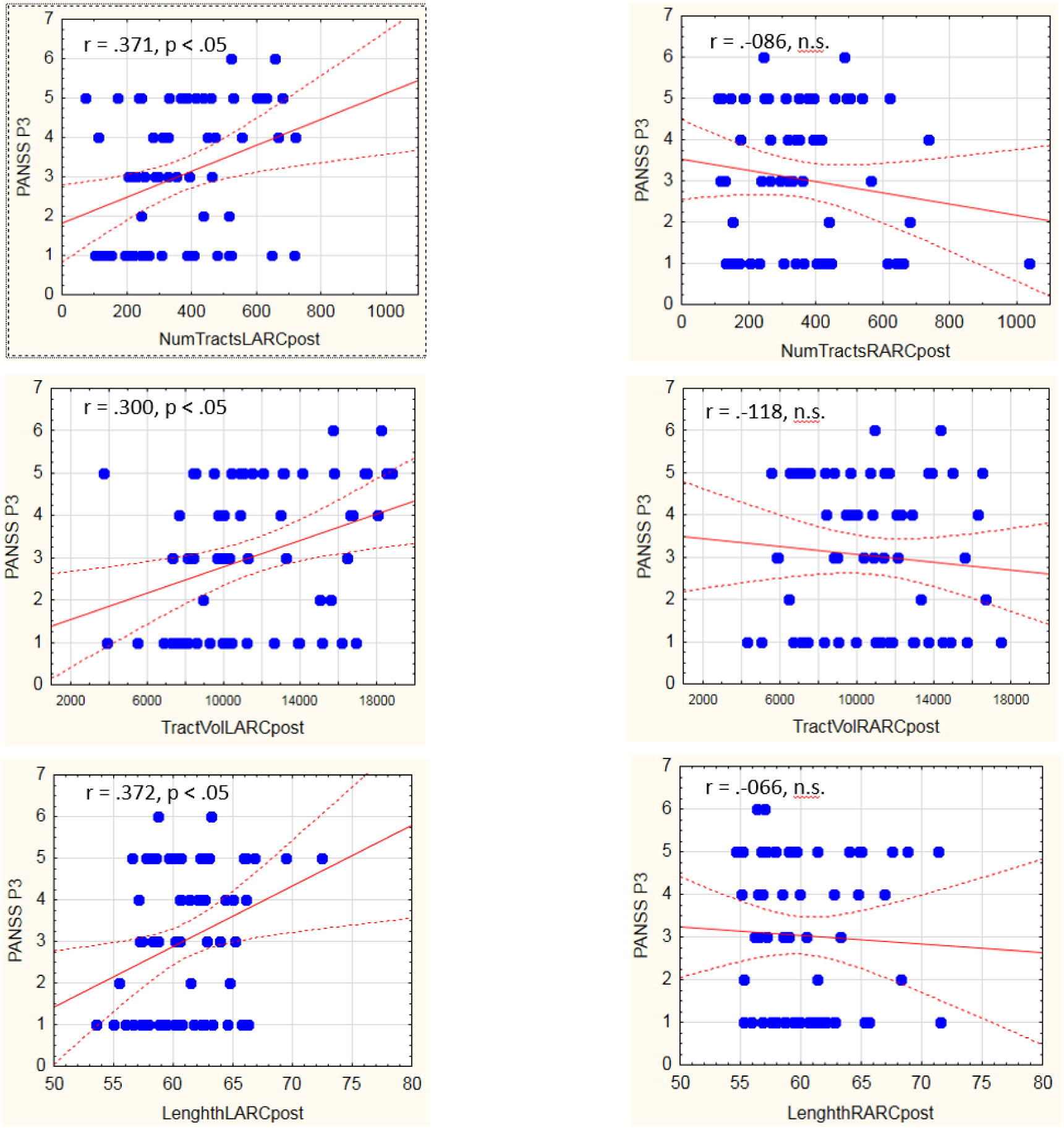
Scatter-plots of the correlation between PANSS P3 scores (y-axis) and number of tracts, tract volume, and tract length for the posterior tract segment, split for the left and right hemisphere. NumTractsLARCpost = Number of tracts, left hemisphere, arcuate fasciculus, posterior segment, TractVolLARCpost = Tract volume, left…., LengthLARCpost = Tract length, left…, and corresponding abbreviations for the right hemisphere (right-hand panel of Figure 4).

## Discussion

The results showed significant effects for the long segments of the arcuate fasciculus (AF) fiber bundle, where patients with severe and frequent hallucinations had longer fibers than healthy controls, and marginally so also when compared to non-hallucinating patients. Interestingly, this was most profound for the left hemisphere arcuate fasciculus bundle, connecting the receptive (Wernicke) and productive (Broca) language areas, respectively. The parameter TractL refers to the average distance the tractography algorithm managed to reconstruct before the tracking abortion threshold (FA criteria or angle) is met. Thus, longer as compared to shorter tracts should make it easier to reconstruct fiber bundles, possibly due to less deterioration by crossing fiber tracks. In the present case, this could suggest that the hallucinating group has increased connectivity between auditory areas in the temporal lobe and frontal and parietal areas associated with language (Fernandez-Miranda et al. 2015). Previous studies have indeed found an increase in white matter volume in the regions connected by the arcuate fasciculus bundle (see e.g. Mitelman et al., 2007), and that greater overall white matter volume predicted higher positive symptom score five years later (Cahn et al., 2006). A second finding in the present study was the significant positive correlations between PANSS P3 scores and left hemisphere AF measures in the posterior segment, largely covering the peri-Sylvian region, including parts of auditory and speech perception cortex. It should be noted that the correlations were found only in the left hemisphere, which is also the language dominant hemisphere (Sperry, 1974; Tervaniemi & Hugdahl, 2003, van der Haegen et al., 2013).

From the present findings it is not possible to disentangle whether it is the larger AF that allows for AVHs to be experienced, or if these segments increase in size due to the presence of AVH. It could indicate that in these individuals, the information flow between language areas could have multiple destinations and origins not found in HC, so that e.g. more tracts reach frontal and parietal areas from the auditory areas, thus leading to a perception of a voice that is not there.

The present white matter tractography results can be seen as extending previous functional studies showing increased BOLD connectivity in hallucinating patients between temporal and frontal language areas (e.g. Diederen et al., 2013; van Lutterwald et al., 2014; Chang et al., 2017). The reason for this suggestion is that an increase in length of the long segment AF fibers could facilitate neuronal connectivity between the areas being connected by the AF in the temporal and frontal lobes, which in turn could contribute to a kind of functional hyper-activity in these regions during hallucinatory episodes (Jardri et al., 2011; Kompus et al., 2011). It has previously been shown that functional hyper-activity in the language areas in the temporal lobe in hallucinating patients (Curcic-Blake et al., 2017; Hugdahl 2015) goes together with reduction in grey matter volume (Neckelmann et al., 2006; Modinos et al., 2013), and that patients with schizophrenia in general have reduced grey matter volume in both temporal and frontal regions (Williams, 2008 for a meta-analysis). The present study extends these previous findings for grey matter abnormality in hallucination-prone individuals, by showing that also white matter structures are affected by hallucinations.

We have focused on tractography and fiber structure in the present study, rather than on the fractional anisotropy (FA) value, which is the more common measure of white matter integrity when using DTI (see Mori, 2007). Although FA reflects axonal diameter, it is primarily measure of the deviation from free diffusion of water molecules, which relates to the restriction of water flow due to fiber myelination (Özarslan et al., 2005). It is therefore interesting to note that there were no significant effects for the FA measure in the present study (cf. Chawla et al., 2019), which could reflect that the critical parameters are fiber length and composition rather than the degree of myelination of the fibers. FA is a measure of water diffusion and is therefore subject to fluctuations across time, while tract parameters, like length, volume and number of tracts, more reflect structural invariants that may better match trait differences in hallucinations as seen in the present study.

In conclusion, we have shown that white matter fibers of the arcuate fasciculus bundle, which connects temporal and frontal language areas in the brain, are affected in patients with severe and frequent hallucinations. Interestingly, increased tract connectivity was observed not only for the long fibers connecting the Wernicke and Broca language areas in the superior temporal and inferior frontal gyri, respectively, but was also observed for the posterior segment fibers, primarily connecting the neurons within the auditory and speech perception areas in the peri-Sylvian region in the left hemisphere. The asymmetry for the long segment fibers should also be noted, there was clearly (see Figure 3) overall longer fibers in the left compared with the right hemisphere, which was seen for both the healthy controls (HC) and with a trend for significance in the frequent hallucinating group (AVH+). Asymmetry of arcuate fasciculus fibers have been observed previously, both in children and adults (Sreedharan et al., 2015; Takao et al., 2011, Ocklenburg et al., 2014), with left hemisphere fibers dominating over right hemisphere fibers. This could be an anatomical correspondence to functional dominance of the left hemisphere for language (Hellige, 1993; van den Noort et al., 2008)). We extend these findings by showing that such an asymmetry also seems to be present in patients with frequent and severe auditory hallucinations, while the asymmetry is absent in schizophrenia patients without hallucinations. This may point to a specific relation between language and the experience of “hearing voices” in a hallucinatory episode, not seen otherwise in schizophrenia patients.

Several limitations should however be noted with the present results. First of all, the sample size was not exceedingly large when splitting the patient group into AVH- and AVH+ sub-groups, which should be kept in mind. Second, we did not find any effects for FA values, which could indicate that the sequence applied was not sensitive enough, or that scanner upgrade could have had an effect, although we could not find evidence for this. It could also be argued that the hallucinatory behavior item of the PANSS interview scale is not unique for auditory verbal hallucinations. Although this is an often claimed argument we do not think it has invalidated our results since there is a clear focus on auditory verbal hallucinations in how the questions and conversations are set up for the PANSS P3 item, and auditory verbal hallucinations is the most frequent mode of hallucinations in schizophrenia. A third limitation to be mentioned is that brain size could have an effect on the tract length parameter, with longer tracts being positively related to larger brains. This is however not a likely explanation for the tract length findings since the age and sex did not differ between groups, which could otherwise have had an effect.

## References

Alderson-Day, B., McCarthy-Jones, S., Fernyhough, C. (2015). Hearing voices in the resting brain: A review of intrinsic functional connectivity research on auditory verbal hallucinations. Neuroscience and Biobehavioral Reviews, 55, 78–87.

Aleman A, Laroi F (2008): Hallucinations: The science of idiosyncratic perception. Washington, DC: American Psychological Association.

American Psychiatric Association (2013). Diagnostic and Statistical Manual of Mental Disorders, 5th Edn. American Psychiatric Association: Washington D.C., USA

Cahn, W, van Haren, NE, Hulshoff Pol, HE, Schnack, HG, Caspers, E, Laponder, DA, Kahn, RS (2006). Brain volume changes in the first year of illness and 5-year outcome of schizophrenia. British Journal of Psychiatry, 189, 381–382.

Catani, M, Craig, MC, Forkel, SJ, Kanaan, R, Picchioni, M, Toulopoulou, T, Shergill, S, Williams, S, Murphy, DG, McGuire, P (2011). Altered integrity of perisylvian language pathways in schizophrenia: relationship to auditory hallucinations. Biological Psychiatry, 70, 1143–1150

Catani, M, Thiebaut de Schotten, M. (2008). A diffusion tensor imaging tractography atlas for virtual in vivo dissections. Cortex, 44, 1105–1132

Chang, X., Collin, G., Xi, Y., Cui, L., Scholtens, L.H., Sommer, I.E., Wang, H., Yin, H., Kahn, R.S., van den Heuvel, M.P. (2017). Resting-state functional connectivity in medication-naïve schizophrenia patients with and without auditory verbal hallucinations: A preliminary report. Schizophrenia Research, 188, 75–81.

Chawla, N., Deep, R, Khnadelwai, S.K., garg, A. (2019). Reduced integrity of superioer longitudinal fasciculus and arcuate fasciculus as a marker of auditory halluicnations in schizophrenia: A DTI tractography study. Asian Journal of Psychiatry, 44, 179 – 186.

Clos M, Diederen KM, Meijering AL, Sommer IE, Eickhoff SB. (2014). Aberrant connectivity of areas for decoding degraded speech in patients with auditory verbal hallucinations. Brain Structure and Function, 219, 581–594.

Curcic-Blake, B., Ford, J.M., Hubl, D., Orlov, N.D., Sommer, I.E., Waters, F., Allen, P., Jardri, R., Woodruff, P.W., David, O., Mulert, C., Woodward, T., Aleman, A. (2017). Interaction of language, auditory and memory brain networks in auditory verbal hallucinations. Schizophrenia Bulletin, 148, 1–20.

Diederen, KM, Neggers, SF, de Weijer, AD, van Lutterveld, R, Daalman, K, Eickhoff, SB, Clos, M, Kahn, RS, Sommer, IE (2013). Aberrant resting-state connectivity in non-psychotic individuals with auditory hallucinations. Psychological Medcine, 43, 1685–1696.

Fernandez-Miranda. J.C., Wang, Y., Pathak, S., Stefaneau, L., Verstynen, L., Yeh, F-C (2015). Asymmetry, connectivity, and segmentation of the arcuate fascicle in the human brain. Brain Structure and Function, 220, 1665–1680.

Ford JM, Morris SE, Hoffman RE, Sommer I, Waters F, McCarthy-Jones S, Thoma RJ, Turner JA, Keedy SK, Badcock JC, Cuthbert BN (2014). Studying hallucinations within the NIMH RDoC framework. Schizophrenia Bulletin 40 (Suppl_4), S295–S304.

Geoffroy, P.A., Houenou, J., Duhamel, A., Amad, A., DeWeijer, A.D., Curcic-Blake, B., Linden, D.E.J., Thomas, P., Jardri, R. (2014). The arcuate fasciculus in auditory-verbal hallucinations: A meta-analysis of diffusion-tensor-imaging studies. Schizophrenia Research, 159, 234–237.

Hellige, J. (1993). Hemispheric asymmetry: What’s right and what’s left. Cambridge, MA: Harvard University Press.

Hubl, D., Koening, T., Strik, W.K., Melie Garcia, L., and Dierks, T. (2007). Competition for neuronal resources: how hallucinations make themselves heard. British Journal of Psychiatry, 190, 57–62.

Hugdahl, K & Sommer, I.E. (2018). Auditory verbal hallucinations in schizophrenia from a levels of explanation perspective. Schizophrenia Bulletin, 44, 234–241

Hugdahl, K. (2017). Auditory hallucinations as translational psychiatry: Evidence from magnetic resonance imaging. Balkan Medical Journal, 34, 504–513

Jardri, R., Pouchet, A., Pins, D., Thomas, P. (2011). Cortical activations during auditory verbal hallucinations in schizophrenia: A coordinate-based meta-analysis. American Journal of Psychiatry, 168, 73–81

Kay, S. R., Fiszbein, A., Opler, L. A. (1987). The positive and negative syndrome scale (PANSS) for schizophrenia. Schizophrenia Bulletin, 13, 261–276.

Kompus, K., Westerhausen, R., Hugdahl, K.. (2011). The “paradoxical” engagement of the primary auditory cortex in patients with auditory verbal hallucinations: A meta-analysis of functional neuroimaging studies, Neuropsychologia, 49, 3361–3369.

Kühn, S., Gallinat, J. (2010). Quantitative meta-analysis on state and trait aspects of auditory verbal hallucinations in schizophrenia. Schizophrenia Bulletin, 38, 779–786.

Leemans A, Jeurissen B, Sijbers J, and Jones DK. ExploreDTI: a graphical toolbox for processing, analyzing, and visualizing diffusion MR data. In: 17th Annual Meeting of International Society for Magnetic Resonance in Medicine, p. 3537, Hawaii, USA, 2009

Looijestijn, J., Diederen, K.L.M., Goekoop, R., Sommer, I.E.C., Daalman, K., Kahn, R.S., Hoek, H.W., Blom, J.D. (2013). The auditory dorsal stream plays a crucial role in projecting hallucinated voices into external space. Schizophrenia Research, 146, 314–319

Mitelman, S.A., Brickman. A.M., Shihabuddin, L., Newmark, R.E., Hazlett, E.A., Haznedar, M., Buchsbaum, M.S. (2007). A comprehensive assessment of gray and white matter volumes and their relationship to outcome and severity in schizophrenia. Neiroimage, 37, 449–462.

Modinos, G., Costafreda, S.G.,van Tol, M-J.,McGuire, P.K, Aleman, A., Allen, P. (2013). Neuroanatomy of auditory verbal hallucinations in schizophrenia: A quantitative meta-analysis of voxel-based morphometry studies. Cortex, 49, 1046–1055

Mori, S. (2007). Introduction to diffusion tensor imaging. Amsterdam, The Netherlands: Elsevier.

Neckelmann, G., Specht, K., Lund, A., Ersland, L., Smievoll, A.I., Neckelmann, D. & Hugdahl, K. (2006). Mr morphometry analysis of grey matter volume reduction in schizophrenia: association with hallucinations. International Journal of Neuroscience, 116, 9–23.

Ocklenburg, S., Hugdahl, K., & Westerhausen, R. (2013). Structural white matter asymmetries in relation to functional asymmetries during speech perception and production. Neuroimagee, 83, 1088–1097.

Özarslan, E. Vemuri, B.C. & Mareci, T. H. (2005). Generalized scalar measures for diffusion MRI using trace, variance, and entropy. Magnetic Resonance in Medicine, 53, 866–876.

Parnas J (2013). On psychosis: Karl Jaspers and beyond. In One Century of Karl Jaspers’ General Psychopathology (Eds. G. Stanghellini, T. Fuch), Oxford University Press: Oxford, UK.), pp. 208–226.

Pienkos, E., Giersch, A., Hansen, M. Humpston, C., McCarthy-Jones, S., Mishara, A., Nelson, B., Park, S., Raballo, A., Sharma, R., Thomas, N., Rosen, C. (2019). Hallucinations beyond voices: A conceptual review of the phenomenology of altered perception in psychosis. Schizophrenia Bulletin, 45, S1, S67–S77.

Sartorius N, Jablensky A, Korten A, Ernberg G, Anker M, Cooper JE, Day R (1986): Early manifestations and first-contact incidence of schizophrenia in different cultures. A preliminary report on the initial evaluation phase of the WHO Collaborative Study on determinants of outcome of severe mental disorders. Psychological Medicine, 16, 909–928.

Scheinost, D., Tokoglu, F., Hampson, M., Hoffman, R., Constable, R.T. (2019). Data-driven analysis of functional connectivity reveals a potential auditory verbal hallucination network. Schizophrenia Bulletin, 45, 415–414.

Shapleske, J., Rossell, S., David, A., McGuire, P., Murray, R.M. (1997). Auditory hallucinations and the temporal cortical response to speech in schizophrenia: A functional magnetic resonance imaging study. American Journal of Psychiatry, 154, 1676–1682

Shreedharan, R.M., Menon, A.C., James, J.S., Kesavadas, C, Thomas, S.V. (2015). Arcuate fasciculus laterality by diffusion tensor imaging correlates with language laterality by functional MRI in preadolescent children. Neuroradiology, 57, 291–297.

Sommer IE, Clos M, Meijering AL, Diederen KM, Eickhoff SB. (2012). Resting state functional connectivity in patients with chronic hallucinations. PLoS One, 7, e43516.

Sperry, R. W. (1974). Lateral specialization in the surgically separated hemispheres, The Neuroscience: Third Study Program (pp. 5–19). Cambridge: MIT Press.

Takao, H., Hayashi, N., Ohtomo, K. (2011). White matter asymmetry in healthy individuals: a diffusion tensor imaging study using tract-based spatial statistics. Neuroscience, 193, 291–299.

Tervaniemi, M. & Hugdahl, K. (2003). Lateralization of auditory-cortex functions. Brain Research Reviews, 43, 231–246

Van der Haegen, L., Westerhausen, R., Hugdahl, K., Brysbaert, M (2013). Speech dominance is a better predictor of functional brain asymmetry than handedness: A combined fMRI word generation and behavioral dichotic listening study. Neuropsychologia, 51, 91–97

Van Lutterveld, R., Diederen, K.M.J., Otte, W.M., Sommer, I.E., (2014). Network analysis of auditory hallucinations in nonpsychotic individuals. Human Brain Mapping, 35, 1436–1445.

Van den Noort, M., Specht, K., Rimol, LM, Ersland, L., & Hugdahl, K. (2008) A new verbal reports fMRI dichotic listening paradigm for studies of hemispheric asymmetry. Neuroimage, 40, 902–911

Waters FA, Badcock JC, Michie PT, Maybery MT (2006): Auditory hallucinations in schizophrenia: Intrusive thoughts and forgotten memories. Cognitive Neuropsychiatry 11, 65–83.

Williams, L. M. (2008). Voxel based morphometry in schizophrenia: Implications for neurodevelopmental connectivity models and cognition and affect. Expert Reviews in Neurotherapeutics, 8, 1029–1036.

Woodruff, P.W.R., Wright, I. Bullmore, E., Brammer, M., Howard, R. J., Williams, S. C.R., Shapleske, J., Rossell, S., David, A., McGuire, P., Murray, R.M. (1997). Auditory hallucinations and the temporal cortical response to speech in schizophrenia: A functional magnetic resonance imaging study. American Journal of Psychiatry, 154, 1676–1682

Zhuo, C., Zhou, C., Lin, X., Tian, H., Wang, L., Chen, C., Ji, F., Xu, Y., Jian, D (2020). Common and distinct global functional connectivity density alterations in drug-naïve patients with first-episode major depressive disorder with and without auditory verbal hallucination. Progress in Neuropsychopharmacology & Biological Psychiatry, 96, 1–6, https://doi.org/10.1016/j.pnpbp.2019.109738

